# Cellular Crowding Influences Extrusion and Proliferation to Facilitate Epithelial Tissue Repair

**DOI:** 10.1101/324301

**Authors:** Jovany Franco, Youmna Atieh, Chase D. Bryan, Kristen M. Kwan, George T. Eisenhoffer

**Affiliations:** Department of Genetics, The University of Texas MD Anderson Cancer Center, Houston, Texas 77030; Department of BioSciences, Rice University, W100 George R. Brown Hall, Houston, TX 77251-1892; Department of Human Genetics, The University of Utah, Salt Lake City, Utah 84112; Genetics and Epigenetics Graduate Program, The University of Texas Graduate School of Biomedical Sciences at Houston, The University of Texas MD Anderson Cancer Center, Houston, Texas 77030

**Author notes:** Corresponding Author and Lead Contact: George T. Eisenhoffer, PhD, Department of Genetics, Unit 1010, The University of Texas MD Anderson Cancer Center, 1515 Holcombe Blvd., Houston, Texas 77030-4009.

## Abstract

Epithelial wound healing requires a complex orchestration of cellular rearrangements and movements to restore tissue architecture and function after injury. While it is well-known that mechanical forces can affect tissue morphogenesis and patterning, how the biophysical cues generated after injury influence cellular behaviors during tissue repair is not well understood. Using time-lapsed confocal imaging of epithelial tissues in living zebrafish larvae, we provide evidence that localized increases in cellular crowding during wound closure promote the extrusion of non-apoptotic cells via mechanically regulated stretch-activated ion channels (SACs). Directed cell migration toward the injury site promoted the rapid changes in cell number and generated shifts in tension at cellular interfaces over long spatial distances. Perturbation of SAC activity resulted in failed extrusion and increased proliferation in crowded areas of the tissue. Together, we conclude that localized cell number plays a key role in dictating cellular behaviors that facilitate wound closure and tissue repair.

## Introduction

Epithelial tissues provide a protective barrier for the body and organs throughout development and adult life. As the first line of defense, these multicellular structures are continually remodeled to sustain tissue form and function. Precise control and coordination of cell division and death is required to retain constant cell numbers during homeostasis and repair the tissue after injury. As individual epithelial cells are physically coupled together, the addition or deletion of cells causes neighboring cells to be stretched, squeezed, move and change shape (Mason and Martin, 2011; Fernandez-Sanchez *et al.*, 2015; Navis and Nelson, 2016). This cellular reorganization exerts mechanical forces on and between cells that are transduced via specific biochemical signals that influence their behavior. Thus, deciphering how mechanical forces coordinate multicellular behaviors is essential to our understanding of epithelial tissue maintenance and repair after injury.

Mechanical forces generated by living cells are crucial for the control of embryological development, morphogenesis and tissue patterning (Mammoto and Ingber, 2010; Davidson, 2011; Heisenberg and Bellaiche, 2013). Cells sense and react to forces imposed by neighbors based on changes in membrane curvature and shearing of the underlying actin cortex (Dreher *et al.*, 2016). Cells can also generate forces by dynamic reorganization of F-actin structures and the associated myosin motors that drive cellular shape changes during tissue morphogenesis (Pasakarnis *et al.*, 2016). The correct timing, location and magnitude of mechanical forces subjected on and between cells are critical for many morphogenetic processes (Lemke and Schnorrer, 2017; Vining and Mooney, 2017), yet the inability to visualize these dynamic events in living epithelial tissues has been an impediment to understanding how biophysical forces can direct epithelial cell function.

Here we use high-resolution time-lapse microscopy of epithelial tissues in living zebrafish larvae to track individual cellular behaviors and population dynamics after injury. Our analysis revealed that localized changes in cell numbers generate mechanical forces that influence the elimination of damaged or excess cells, or production of new cells. Localized increases in cellular crowding promoted the extrusion of non-apoptotic cells from the wound edge via mechanically regulated stretch-activated ion channels (SACs), key mediators of mechanosensitivity (Coste *et al.*, 2010; Sachs, 2010; Olsen *et al.*, 2011). Interestingly, perturbation of SAC activity resulted in failed cellular extrusion and increased proliferation in crowded areas of the tissue. Together, our data establish a temporal series of mechanically-induced cellular and molecular events that facilitate wound closure and tissue repair.

## Results and Discussion

To visualize the cytoskeleton in living epithelial cells during epithelial tissue maintenance and repair after injury *in vivo* and in real time, we imaged transgenic *Et(Gal4-VP16)^zc1044A^; Tg(UAS:Lifeact-GFP)* zebrafish larvae with F-actin fluorescently labeled in surface epithelial cells under homeostatic conditions and immediately after amputation (Figure 1, A-B, and Supplemental Figure S1A). Under homeostatic conditions, epithelial tissues in non-amputated 4 day post-fertilization (dpf) larvae eliminate damaged or excess cells by extrusion (Figure 1A and Supplemental Video S1), as identified by the formation of an actomyosin ring in neighboring cells that contracts and ejects the cell from the tissue (Rosenblatt *et al.*, 2001). We found the number of extruding cells significantly increased after injury from 0.33 ± 0.19 cells under homeostatic conditions to 6.38 ± 0.60 cells at 15 minutes post-amputation (mpa), and continued at similar levels for the next 45 minutes (Figure 1, B-C, and Supplemental Video S2). Further analysis with cell type specific markers revealed that extruded cells originate from both the apical periderm layer, as well as the basal layer of p63-positive progenitor/stem cells (Supplemental Figure S1, B-C).

**Figure 1.**
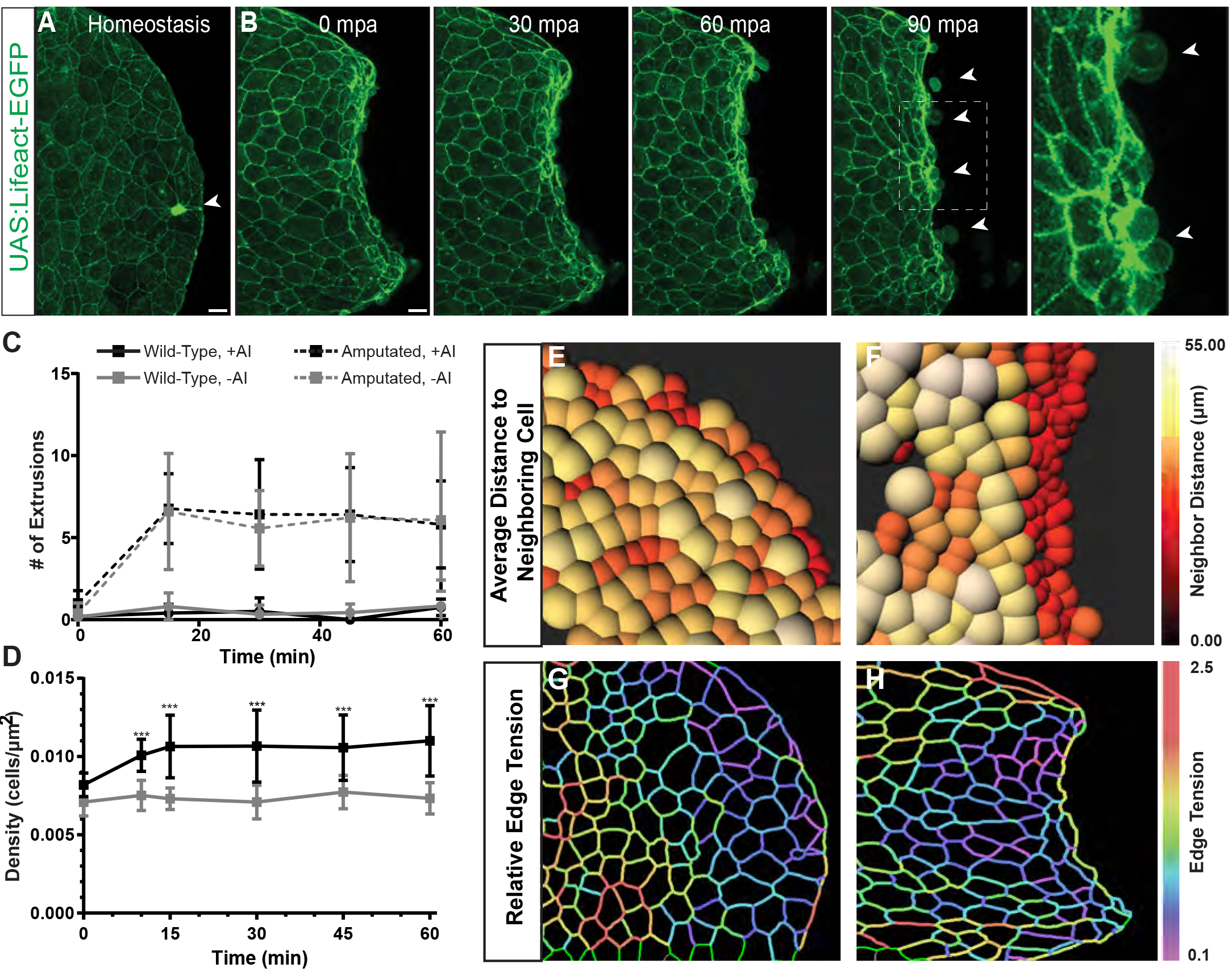
Increased cell density at the wound site promotes cell extrusion. (A-B) Max intensity projection still images of confocal live imaging of (A) non-amputated and (B) amputated *Et(Gal4-VP16)zc1044A; Tg(UAS:LifeAct-GFP)* 4 day post-fertilization (dpf) zebrafish larvae acquired every 15 min. Arrowheads indicate extrusions. Scale bar, 25 μm. (See Supplemental Video S1) (C) Quantification of cell extrusions after amputation in untreated (n=72) and apoptosis inhibitor-treated (n=56) larvae over time, which were significantly increased compared to non-amputated controls both untreated (n=28) and treated (n=26). (D) Quantification of cell density from maximum intensity projections from confocal images of fixed amputated embryos (n=132) and non-amputated controls (n=62) over time. (E-F) Density maps of unamputated (E) and amputated (F) DAPI stained larvae at 30 minutes post amputation (mpa). Color-coded spheres denote distance to the nearest neighboring cell. (G-H) Tension maps from confocal images of non-amputated (G) and amputated (H) *Et(Gal4-VP16)zc1044A; Tg(UAS:LifeAct-GFP)* larvae at 30mpa using CellFit. Color code shows the inferred relative tensions at the cell edges ranging from 0.1 to 2.5.

The extrusion pathway can operate in two distinct manners: one that is stimulated by cellular crowding and the other induced by caspase-activation to remove apoptotic/damaged cells (Eisenhoffer *et al.*, 2012). To determine whether the cells being eliminated by extrusion were undergoing apoptosis, we examined cleaved caspase-3 activity after amputation. While we observed increased levels of activated caspase-3 in larvae treated with UV-C to induce DNA damage, activated caspase-3 was rarely detected in extruding cells (0.27%, 2/742 cells) near the wound edge (Supplemental Figure S2, A-E). Moreover, the number of extrusion events after amputation was also unchanged in amputated larvae treated with a pharmacological inhibitor of apoptosis (Apoptosis Inhibitor II, NS3694) (Gauron *et al.*, 2013) when compared to untreated embryos (Figure 1C). We conclude from this data that non-apoptotic cells are cleared from the wound site by extrusion after amputation.

Cellular crowding plays a key role in promoting extrusion of non-apoptotic cells (Eisenhoffer *et al.*, 2012; Marinari *et al.*, 2012; Eisenhoffer and Rosenblatt, 2013), and as a result, we quantified the distance between all nuclei under homeostatic conditions and at different times after amputation. This approach showed distinct areas of localized cellular crowding at the fin edge under homeostatic conditions (Figure 1E), and uncovered a significantly increased number of cells per area (76.97%) near the injury site (<100 μm) within 10 mpa (Figure 1, D-F and Supplemental Video S3). The cell density increase occurs 5 minutes before the observed extrusion events (Figure 1, C-D and Supplemental Video S2), suggesting a temporal link between crowding and non-apoptotic cell extrusion. Rapid increases in cell density can generate mechanical forces to influence a range of biological processes, including cell extrusion, and as a result we sought to define the contribution of cellular crowding and mechanical forces at the wound site.

Cellular shape and size is a reflection of the forces that epithelial cells experience during the regeneration process (Chanet and Martin, 2014; Aragona *et al.*, 2017). To define the forces exerted on epithelial cells under homeostatic conditions and after amputation, we analyzed our images using CellFIT, the Cellular Force Inference Toolkit, which assumes that cell shapes are determined by interfacial tensions at cell edges (Brodland *et al.*, 2014). Tensions at the interfaces between cells under homeostatic conditions varied depending on the location of the cell within homeostatic epithelial tissue. Tensions of cells over 100 μm away from the edge were the highest and gradually decreased towards the fin edge, pointing towards a more active flow of cell migration in the anterior part of the developing tail fin (Figure 1G and Supplemental Figure S2F). After injury, edge tension was also significantly increased in cells located more anterior from the amputation site, while tension in cells in the middle section significantly decreased (Figure 1H and Supplemental Figure S2F). This shift in tension suggests an active migratory front of cells that induces crowding and subsequent extrusion towards the edge. Together, this approach allowed us to reveal areas of the tissue likely to experience the greatest shifts in tensions, as well as predict areas of active migration during wound repair.

To capture the tissue-wide dynamics and individual cellular behaviors that contribute to mechanical forces during wound closure and repair, we examined both crowding and extrusion in real time by imaging *Tg(Ubi:H2A-EGFP-2A-mCherry-CAAX)* larvae with all nuclei and cell membranes fluorescently labeled and tracking individual cells over time after amputation. To directly test if localized cellular crowding promotes extrusion, we quantified the distance to the nearest neighbor in individual cells that would go on to extrude and found the likelihood of a cell being extruded dramatically increased as the cell density increases by more than a factor of 1.4 (Figure 2, A-C). These results are consistent with the 1.4-1.6x critical threshold range previously reported using *in vitro* crowding studies (Eisenhoffer and Rosenblatt, 2013) and mathematical modeling (Shraiman, 2005), and supports the idea of a critical crowding concentration that activates extrusion of non-apoptotic cells in living tissue. Tracking groups and individual cells over the first 30 minutes after amputation, we found cells within the tail fin exhibited a 3.52 fold increase in average distance moved, and with a 1.7 fold increase in average speed, when compared to similar regions in homeostatic controls (Figure 2D and Supplemental Figure S3, A-G). Importantly, this analysis revealed that speed and displacement varied depending on the distance from the wound site. Cells further from the injury site move with increased speed and show a greater displacement compared to those closest to the wound edge, 53% and 21% (n = 36), respectively (Figure 2D and Supplemental Figure S3, A-E). Overlaying these live imaging data on our force inference map predicts the cells undergoing the most movement experience elevated tension at cellular interfaces, and the collective displacement of cells increases crowding at the wound site.

**Figure 2.**
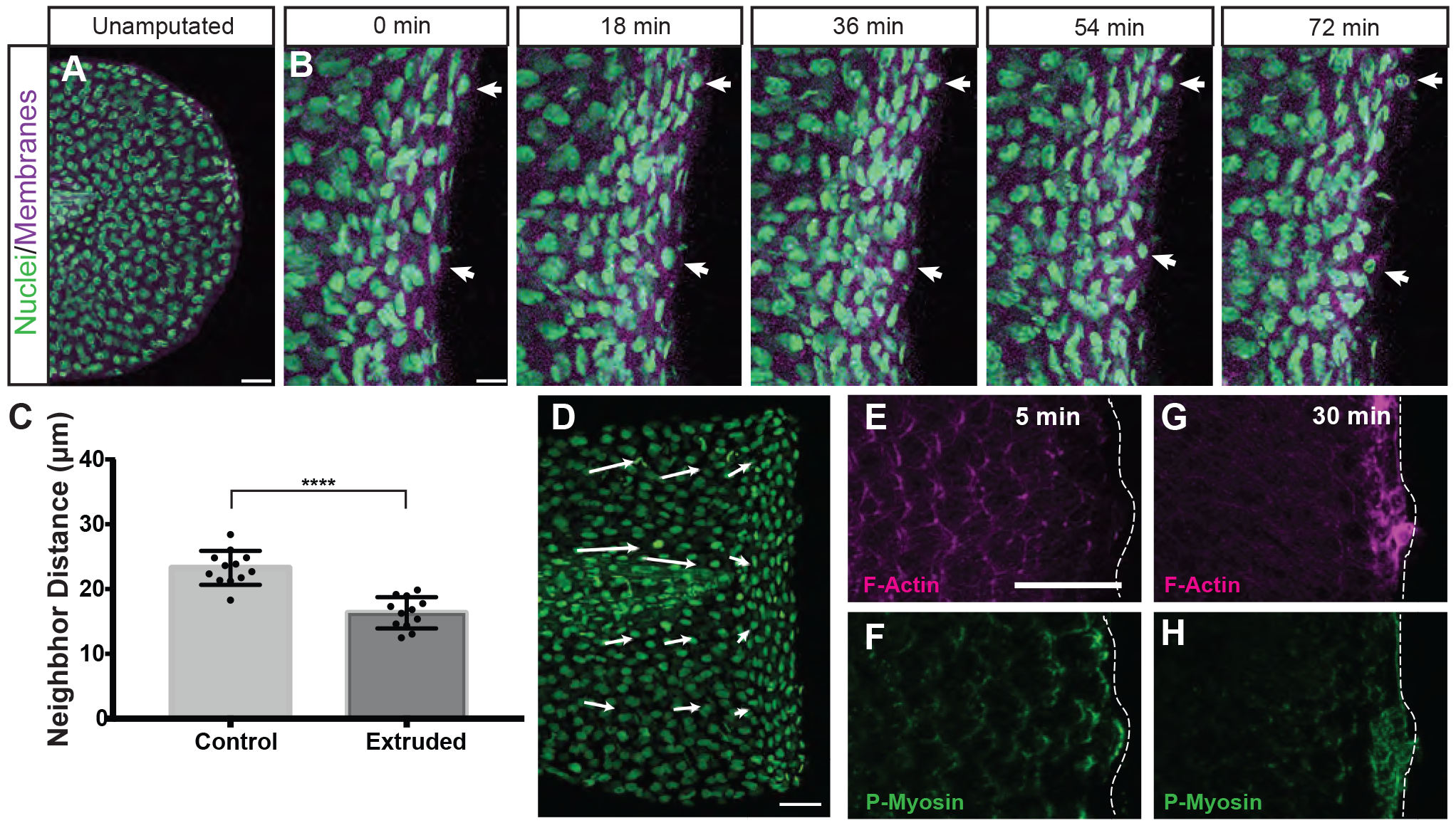
Visualizing localized crowding that promotes cell extrusion in living tissue. (A-B) Max intensity projection still images of confocal live imaging of (A) non-amputated and (B) amputated *Tg(Ubi:H2A-EGFP-2A-mCherry-CAAX)* larvae acquired every 6 minutes. Arrowheads indicate cells during the process of extrusion. Scale bars, (A) 50 μm and (B) 20 μm. (C) Quantification of distance to nearest neighbor during the process of extrusion. Mean values are plotted from 23 extruding cells and 23 non-extruding cells analyzed from three independent datasets. (D) Vector map of the trajectory of displacement of groups of cells in the respective regions. The length and direction of arrow represents quantification of cellular displacement based on tracking from time-lapse imaging in amputated embryos (n=72 cells, 6 independent datasets) compared to non-amputated controls (n=36 cells, 3 independent datasets) over 60 minutes post-amputation (mpa). Scale bar, 50 μm (E-H) Max intensity projection images of fixed embryos stained for F-actin (E,G) and p-myosin (F,H) at 5 and 30 mpa. Scale bar, 50 μm.

Cell movement involves tight regulation of F-actin-binding and non-muscle myosin II activity to control membrane tension and protrusion necessary for migration (Watanabe *et al.*, 2007; Chen *et al.*, 2014). Actomyosin contractions are also thought to be major contributors to the tensions observed at the cell-cell interfaces (Fernandez-Gonzalez and Zallen, 2009; Mason and Martin, 2011; Brodland *et al.*, 2014). Consequently, we next examined F-actin filaments and non-muscle myosin II in epithelial cells within regions of the injured tissue predicted to have differential tension. Phosphorylated-myosin light chain II was localized to cells near the amputation site within 5 mpa, and interestingly, was significantly enriched on the edges of cells facing the wound site (Figure 2, E-H and Supplemental Figure S3, H-I). We also observed differential myosin localization in epithelial cells undergoing cytoskeletal reorganization during extrusion at 30 mpa (Figure 2, E-H). Together, these data support the idea that mechanical forces regulate myosin-mediated cytoskeletal reorganization and cell movement as part of wound closure (Kobb *et al.*, 2017) and further suggest that the resulting localized changes in density contribute to the elimination of excess cells.

These data promoted us to define how shifts in cell-and tissue-level forces after injury influence cellular events in specific locations of the tissue. To disrupt the ability of cells to detect and respond to crowding or tension-induced events, we compared wound closure and crowding-induced extrusion after injury following alteration of stretch-activated ion channels (SACs), key mechano-regulators of crowding-induced non-apoptotic extrusion (Eisenhoffer *et al.*, 2012; Kim *et al.*, 2015). After 60 minutes of observation, an average of 23 extrusion events had occurred in nearly all of the control amputated larvae, whereas treatment with gadolinium trivalent cations (Gd3+) to perturb SACs resulted in a failure of cells to complete extrusion and detach from tissue over time (Figure 3, A-E). Failure to complete the extrusion process resulted in accumulation of undetached cells at the amputation site and a significant increase in width of the wound, from 1.1 ± 0.5 μm to 8.8 ± 1.5 μm, when compared to untreated injured larvae (Figure 3, C’-F). Collectively, our results suggest that crowding-induced non-apoptotic cell extrusion occurs at the wound site and is regulated by mechanosensitive SACs.

**Figure 3.**
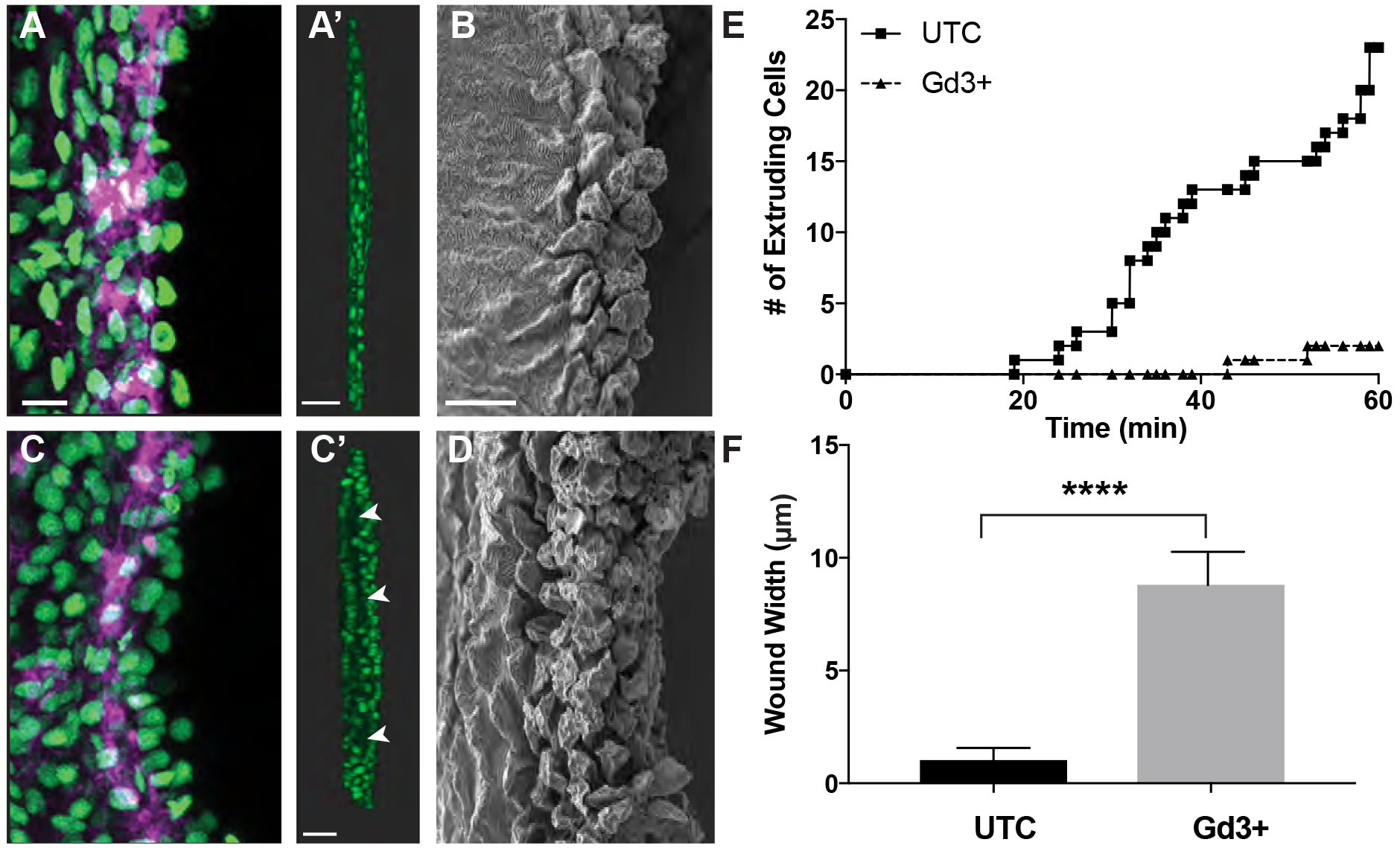
Blocking stretch-activated ion channels (SACs) causes failed extrusion. (A-D) Max projection confocal images and of the wound site at 30 mpa in untreated (A) or Gd3+ treated (C) amputated *Tg(Ubi:H2A-EGFP-2A-mCherry-CAAX)* larvae (Scale bar, 10 μm), and rotated images of the wound site (A’, C’). Scale bar, 50 μm. Arrowheads to denote gaps and areas of the wound that have not fully closed. SEM images (B, D) of the wound site at 30 mpa in untreated or Gd3+ treated amputated larvae. Scale bar, 10 μm. (E) Quantification of successful extrusion events in untreated and Gd3+ treated amputated *Tg(Ubi:H2A-EGFP-2A-mCherry-CAAX)* larvae over time. (F) Quantification of the width of the wound site.

Tightly controlled proliferation of progenitor cells upon amputation is a hallmark of epimorphic regeneration (Kawakami *et al.*, 2004; Rojas-Munoz *et al.*, 2009). To determine how mechano-sensation of cellular crowding could impact the proliferative response required for successful regeneration, we analyzed the number of cells actively synthesizing DNA, as measured by BrdU incorporation, at different locations within the tissue 3 hours after injury. Amputated larvae showed a dramatic increase in the number of proliferating cells compared to homeostatic controls, and interestingly, the proliferating cells were located in areas of predicted elevated edge tension, >100um back from the wound edge (Figure 4, A-C). This finding is in agreement with the notion that cell stretching induces proliferation (Gudipaty *et al.*, 2017). Conversely, inhibiting the cells’ capacity to sense mechanical forces by altering SAC activity by Gd3+ treatment led to proliferation in areas of increased cell density, closer to the wound site, (Figure 4, C-F). These data suggest that the ability to detect and respond to crowding-induced mechanical forces at the wound site is required for both the elimination of non-apoptotic cells by extrusion and spatial and temporal control of proliferation to replace the cell types lost to injury.

**Figure 4.**
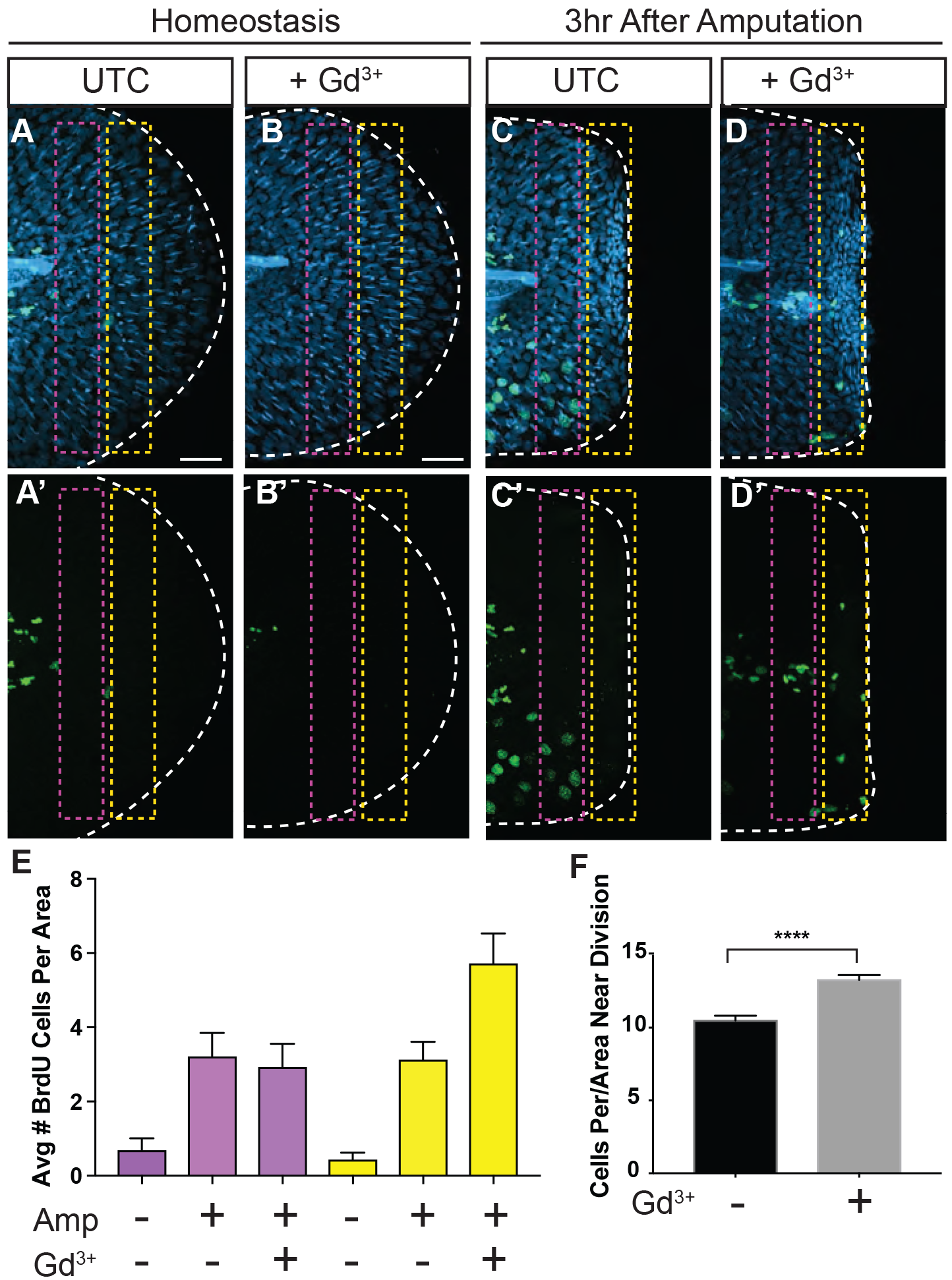
Proliferation in crowded areas after inhibition of SACs during regeneration. (A-D) Max intensity projection images of BrdU incorporation (green-dividing cells) during homeostasis and at 3hr after amputation in fixed Gd3+ treated and untreated DAPI-stained (blue-nuclei) larvae. Scale bar, 50μm. (E) Quantification of the distance of BrdU positive cells from the wound edge. (F) Cell density (cells/μm2) near the proliferating cells in untreated and Gd3+ treated larvae.

In conclusion, our high resolution *in-vivo* imaging based approach has yielded a detailed map of cellular events that generate and result from shifts in mechanical forces within a living epithelial tissue after injury. The generation and transmission of mechanical forces can influence cell behaviors associated with wound repair (Zulueta-Coarasa and Fernandez-Gonzalez, 2017), yet how these events are coordinated in space and time is not well understood. The intercalation and exchange of neighbors, orthogonal cell displacement, collective cell migration and crowding-induced epithelial cell extrusions promoting wound closure and repair in the larval zebrafish fin after injury is similar to that seen after fusion of the mammalian secondary palate during development (Kim *et al.*, 2015) and after fin amputation in adult zebrafish (Chen *et al.*, 2016). The analyses presented here identify a critical threshold of crowding that promotes extrusion in living epithelia that is consistent with previous reports for crowding-induced extrusion and delamination (Shraiman, 2005; Marinari *et al.*, 2012). This is supported by the observation that perturbation of SACs causes defective non-apoptotic extrusion and cell accumulation (Figure 3, A-F) (Eisenhoffer *et al.*, 2012; Kim *et al.*, 2015), as well as aberrant proliferation in crowded areas (Figure 4, C’-F). Our data support the conclusion that the inability of cells to sense changes in mechanical forces may lead to the breakdown of both contact inhibition of locomotion (Abercrombie and Heaysman, 1954; Stramer and Mayor, 2016) and growth (Stoker and Rubin, 1967; Abercrombie, 1979; McClatchey and Yap, 2012), hallmarks of oncogenic cells. Together, our results highlight an essential role for mechanical forces in coordinating multi-cellular behaviors during epithelial tissue maintenance and regeneration after injury.

## Author Contributions

G.T.E designed the research. J.J.F and Y.M.A performed experiments and analyzed data. C.D.B and K.M.K generated key reagents and assisted in interpretation of the resulting data. J.J.F and G.T.E wrote the paper, and all authors provided edits.

## Acknowledgements

We thank members of the MD Anderson Genetics Department, the Texas Medical Center Aquatics community, and the Galko and Qutub laboratories for scientific discussions, suggestions, and comments. This work was supported by the Cancer Prevention Institute of Texas, RR14007, to GTE. KMK was supported by the NEI/NIH (R01 EY025378, R01 EY025780). CDB was supported by the University of Utah Developmental Biology Training Grant (NIH T32HD007491). We thank Kenneth Dunner Jr. at the UT MD Anderson High Resolution Electron Microscopy Facility for help with the electron microscopy data. The High Resolution Electron Microscopy facility was supported by CCSG grant NIH P30CA016672.

## Materials and Methods

Further information and requests for resources and reagents should be directed to and will be fulfilled by the Lead Contact, George T. Eisenhoffer.

### Zebrafish and Wound Healing Assay

Zebrafish maintained under standard laboratory conditions with 14 hr light, 10 hrs dark cycle. Embryos kept in E3 embryo medium at 28.5°C and staged as described in (Kimmel *et al.*, 1995). For the wound closure and repair assay, four day post-fertilization (dpf) larvae were anesthetized with 0.04% tricaine and amputated ~100μm (98.4μm average, n=285) posterior of the notochord using a scalpel. Assays were performed in the wild-type AB* background. The zebrafish used in this study were handled in accordance with the guidelines of the University of Texas MD Anderson Cancer Center Institutional Animal Care and Use Committee.

### Transgenic zebrafish lines

The GAL4 enhancer trap line *Et(Gal4-VP16)^zc1044A^* was used to drive expression of *Tg(UAS-E1b:nsfB-mCherry)* or *Tg(UAS-E1b:Lifeact-EGFP)^zr2^.* The *Et(Gal4-VP16)^zc1036a^;Tg(UAS-E1b:nsfB-mCherry)* line was also used in combination with *Tg(p63:EGFP)* (Eisenhoffer *et al.*, 2017).

*Transgenic constructs and establishment of stable lines Tg(Ubi:H2A-EGFP-2A-mCherry-CAAX)^z201^* was generated using the Tol2kit (Kwan et al., 2007). A 5’ Gateway entry vector containing the zebrafish *ubiquitin* promoter (Mosimann et al., 2011) was combined with a middle entry clone of H2A-EGFP (pME-H2A-EGFP [no stop codon]) and a 3’ entry clone containing a viral 2A peptide (Provost et al., 2007) upstream of mCherry-CAAX and an SV40 late polyA signal sequence (p3E-2A-mCherry-CAAX-pA). These three pieces were recombined to generate the full transgene expression construct in the pDestTol2pA3 destination vector, which contains I-SceI meganuclease sites within the miniTol2 ends. The complete expression plasmid was coinjected with *Tol2* transposase (30 pg DNA + 25 pg transposase RNA) into the cell of a 1-cell embryo. Alternatively, the complete expression plasmid was coinjected with I-SceI meganuclease (30 pg DNA + 2.5 units I-SceI) into the cell of a 1-cell embryo. Injected embryos were screened for fluorescence reporter expression at 24 hpf and raised to adulthood.

### Pharmacological treatments and UV exposure during the wound healing assay

For apoptosis inhibitor assays, embryos treated 18 hrs prior to amputation and during recovery with 10μM apoptosis inhibitor NS3694 (Santa Cruz Biotech) in DMSO diluted in E3. For gadolinium assays, embryos treated with 100 μM gadolinium(III) chloride hexahydrate (Sigma Aldrich) in E3 for 3 hours prior to amputation and during recovery, up to one hour. For UV assays, embryos exposed to 2 min UV-C, 4 hours before amputation.

### Fixation and Immunofluorescence of Zebrafish Larvae

Zebrafish embryos fixed overnight at 4°C with 4% paraformaldehyde in PBS, 0.05% TritonX-100 (PBSTx 0.05%), washed 15 min in PBSTx 0.5%, blocked 2 hrs at room temperature in block buffer (PBS, 1% DMSO, 2 mg/mL bovine serum albumin (BSA), 0.5% Triton X-100, 10% heat inactivated goat serum). Larvae then incubated overnight at 4°C in primary antibodies (below) in block buffer. Samples were washed for 2 hrs in PBSTX 0.5%, incubated in block solution for 2 hrs at room temperature, and then incubated overnight at 4°C in secondary antibodies (below) in block solution. Specimens were then washed 1 hour in PBSTx 0.5%, incubated in DAPI (1:1000) in PBSTx 0.5% for 30 minutes, rinsed with PBS, and the tail portion of the larvae was mounted on sealed glass slides containing 80% glycerol.

### BrdU incorporation and detection

For proliferation assays, larvae were soaked in 10 mM BrdU (Sigma Aldrich), E3+5% DMSO for 1 hr, followed by 30 min recovery and fixation. Larvae washed 15 min in PBSTx 0.5%, 20 min in ddH_2_O, 45 min in 2N HCl in ddH_2_O+0.5% Triton-X, washed 15 min in PBSTx 0.5%, and detection was performed according to protocol above.

### Primary Antibodies

Rat anti-BrdU (Abcam, 1:200), Rabbit anti-Cdh1 (GeneTex, 1:100), Rabbit anti-cleaved Caspase-3 (BD Pharmingen, 1:700), Rabbit anti-phospho-Myl9 (Thr18, Ser19) (Invitrogen, 1:200)

### Secondary Antibodies

Goat anti-Rabbit Alexa Fluor 488 (Invitrogen, 1:200), Donkey anti-Rabbit Alexa Fluor 568 (Invitrogen, 1:200), Goat anti-Rat Alexa Fluor 488 (Invitrogen, 1:200), Phalloidin Alexa Fluor 647 (Invitrogen, 1:50)

### Image Acquisition and Time-lapse confocal microscopy

All confocal images acquired as z-stacks using 20x objective on a Zeiss LSM 800 Confocal Microscope and Zen 2 software. Time-lapse images acquired by anesthetizing day 4 embryos with 0.04% tricaine in E3 and mounting in 1% low melt agarose in 10mm MatTek culture dish. Embryos imaged starting at 5 mpa. Images presented as maximum intensity projections made on Zen 2 software.

### Image Quantification

All quantifications were conducted using maximum intensity projections. Extruding cells were defined by presence of a multi-cellular actin ring and defined nuclei above the actin ring. Apoptotic cells quantified by manually counting activated caspase-3 positive cells posterior to notochord. Cell density was calculated by manually counting DAPI-stained nuclei located posterior to the notochord and defining the area using contour tool in Zen 2. Quantification of crowding near BrdU positive cells was conducted by defining a square of 1143 μm^2^ centered around the BrdU positive cell nuclei and counting DAPI-stained nuclei within the defined region. Localization of anti-phospho-Myl9 was measured by using contour tool to trace around cell membranes and measure fluorescent intensity of membranes for the anterior half and posterior half of the cell, where polarization = anterior intensity ÷ posterior intensity.

### Image processing and cell tracking using Imaris

Z-stack rotations done using Bitplain Imaris x64 9.0.2 using either Maximum Intensity Projection or Blend display mode. Cell migration measurements for *Tg(UAS:LifeAct-GFP)* larvae acquired using Imaris with “Cell” function to detect membranes using the following settings: detection type: membrane; filter type: local contrast; cell tracking algorithm: autoregressive motion; cell smallest diameter: 12.5 μm; membrane detail: 1.25 μm; intensity ≥ 40 μm; quality ≥ 0.93. Measurements for *Tg(Ubi:H2A-EGFP-2A-mCherry-CAAX)* larvae acquired using Imaris with “Spots” function to detect nuclei using the following settings: estimated diameter: 11 μm; quality: >21; tracking algorithm: autoregressive motion; max gap size: 3. Values for instantaneous speed were calculated for each frame within first 30 mpa, along with values for displacement after 1 hpa, for each cell tracked. Nearest neighbor analysis conducted using Imaris to detect cells as above with “Measuring Points” function to measure distance between center of cell of interest and neighbors at frame where extrusion begins. Threshold calculated by dividing average nearest neighbor distance of control cells by average nearest neighbor distance of extruded cells. Data exported and analyzed using GraphPad Prism 7.0.

### Inference of tension using Cell Fit

For tension measurements, image segmentation was performed using an automated ImageJ plugin *Tissue Analyzer*. Tension maps were generated on segmented images using the CellFIT analysis software. CellFIT formulation was applied to the entire mesh using nearest segment tangent vectors.

### Statistical Analysis

All statistical analyses conducted using unpaired Welch’s t-tests. *: p < 0.05, **: p < 0.01, ***: p < 0.001, ****: p < 0.0001 using Prism.

### Scanning Electron Microscopy

Samples were fixed in 3% glutaraldehyde + 2% paraformaldehyde in 0.1M cacodylate buffer (pH 7.3). Samples were washed with 0.1M cacodylate buffer (pH 7.3), post-fixed with 1% cacodylate buffered osmium tetroxide, washed with 0.1M cacodylate buffer, and then in distilled water. Samples were treated with Millipore-filtered 1% aqueous tannic acid, washed in distilled water, treated with Millipore filtered 1% aqueous uranyl acetate, and then rinsed with distilled water. They were dehydrated with a series of increasing concentrations of ethanol and transferred to increasing concentrations of hexamethyldisilazane (HMDS) and air dried overnight. Samples were mounted onto double-stick carbon tabs (Ted Pella. Inc., Redding, CA), and mounted onto glass microscope slides. Samples were coated under vacuum using a Balzer MED 010 evaporator (Technotrade International, Manchester, NH) with platinum alloy for a thickness of 25nm and then flash carbon coated under vacuum. Samples were then transferred to a desiccator until examination and imaging in a JSM-5910 scanning electron microscope (JEOL, USA, Inc., Peabody, MA) at an accelerating voltage of 5kV.

## Supplemental Video Files

**Supplemental Video S1. Epithelial Cell Extrusion Under Homeostatic Conditions.** Maximum intensity confocal projection images were acquired 3 minutes and 3 seconds for 54 minutes and 54 seconds *Et(Gal4-VP16)^zc1044a^;Tg(UAS-E1b:Lifeact-EGFP)* under homeostatic conditions. Scale bar, 50 μm.

**Supplemental Video S2. Visualizing early cellular events after injury in a living epithelial tissue.** Maximum intensity confocal projection images acquired every 15 minutes for 1.5 hours of *Et(Gal4-VP16)^zc1044a^;Tg(UAS-E1b:Lifeact-EGFP)* after amputation. Scale bar, 50 μm.

**Supplemental Video S3. Cellular crowding promotes extrusion at the wound site after injury.** Maximum intensity confocal projection of time-lapse imaging of *Tg(Ubi:H2A-EGFP-2A-mCherry-CAAX)* after amputation. Images were acquired every 5 minutes and 51 seconds for 1.5 hours. Scale bar, 50 μm.

